# Ultra-high sensitivity mass spectrometry quantifies single-cell proteome changes upon perturbation

**DOI:** 10.1101/2020.12.22.423933

**Authors:** Andreas-David Brunner, Marvin Thielert, Catherine G. Vasilopoulou, Constantin Ammar, Fabian Coscia, Andreas Mund, Ole B. Hoerning, Nicolai Bache, Amalia Apalategui, Markus Lubeck, Sabrina Richter, David S. Fischer, Oliver Raether, Melvin A. Park, Florian Meier, Fabian J. Theis, Matthias Mann

**Affiliations:** Proteomics and Signal Transduction, Max-Planck Institute of Biochemistry, Am Klopferspitz 18, 82152 Martinsried, Germany; NNF Center for Protein Research, Faculty of Health and Medical Sciences, University of Copenhagen, Blegdamsvej 3B, 2200 Copenhagen, Denmark; EvoSep Biosystems, Thriges Pl. 6, 5000 Odense, Denmark; Bruker Daltonik GmbH, Fahrenheitstraße 4, 28359 Bremen, Germany; Bruker Daltonics Inc., 40 Manning Road, Billerica, MA 01821, United States of America; Helmholtz Zentrum München – German Research Center for Environmental Health, Institute of Computational Biology, Ingolstädter Landstraße 1, Neuherberg, Germany; TUM School of Life Sciences Weihenstephan, Technical University of Munich, Alte Akademie 8, 85354 Freising, Germany; Functional Proteomics, Jena University Hospital, Am Klinikum 1, 07747 Jena, Germany

## Abstract

Single-cell technologies are revolutionizing biology but are today mainly limited to imaging and deep sequencing^1–3^. However, proteins are the main drivers of cellular function and in-depth characterization of individual cells by mass spectrometry (MS)-based proteomics would thus be highly valuable and complementary^4,5^. Chemical labeling-based single-cell approaches introduce hundreds of cells into the MS, but direct analysis of single cells has not yet reached the necessary sensitivity, robustness and quantitative accuracy to answer biological questions^6,7^. Here, we develop a robust workflow combining miniaturized sample preparation, very low flow-rate chromatography and a novel trapped ion mobility mass spectrometer, resulting in a more than ten-fold improved sensitivity. We accurately and robustly quantify proteomes and their changes in single, FACS-isolated cells. Arresting cells at defined stages of the cell cycle by drug treatment retrieves expected key regulators such as CDK2NA, the E2 ubiquitin ligase UBE2S, DNA topoisomerases TOP2A/B and the chromatin regulator HMGA1. Furthermore, it highlights potential novel ones and allows cell phase prediction. Comparing the variability in more than 430 single-cell proteomes to transcriptome data revealed a stable core proteome despite perturbation, while the transcriptome appears volatile. This emphasizes substantial regulation of translation and sets the stage for its elucidation at the single cell level. Our technology can readily be applied to ultra-high sensitivity analyses of tissue material^8^, posttranslational modifications and small molecule studies to gain unprecedented insights into cellular heterogeneity in health and disease.

## Main

In single-cell analysis, biological variability can directly be attributed to individual cells instead of being averaged over an ensemble or complex tissue^9^. While microscopy has always been single-cell based, specialized deep sequencing technologies have achieved this for systems biological approaches^10–12^. At the level of proteins, the functional actors of cells, single cells are currently studied by antibody-based technologies, which are by necessity directed against previously chosen targets^13–15^. In contrast, mass spectrometry (MS)-based proteomics is unbiased in the sense that it measures all proteins within its range of detection^4,16^. Thus, it would be highly desirable to apply this technology to single cells if the required sensitivity and robustness could be achieved. Previous approaches that employed chemical multiplexing of peptides have labeled a small number of single cells but combined them with a dominant booster channel for MS-analysis^17,18^, which can hamper signal deconvolution^6,19^. Alternatively, proof of principle has been demonstrated for unlabeled approaches using sophisticated sample preparation methods in pico-liter devices^7,20,21^. However, a technology that provides quantitatively accurate MS proteomics data from true single-cells (T-SCP) and answers biological questions is still outstanding.

### Noise-reduced quantitative mass spectra

We recently introduced parallel accumulation – serial fragmentation (PASEF), a mass spectrometric acquisition scheme in which peptide ions are released from a trapped ion mobility (TIMS) device into the vacuum system in concentrated packages^22,23^. Chemical noise is widely distributed as a result of its heterogeneous nature and the ten-fold increased peak capacity due to TIMS (**Fig. 1a, b**)^24^. These precursors can be fragmented in a highly sensitive manner, either in data dependent (ddaPASEF) or data independent (diaPASEF) mode, resulting in very high ion utilization and data completeness^25^. To explore sensitivity limits, we measured a dilution series of HeLa cell lysate from 25 ng down to the equivalent of a few single cells on a quadrupole time-of-flight instrument (TIMS-qTOF). This identified more than 550 proteins from 0.8 ng HeLa lysate with the dda acquisition mode and a conservative MaxQuant analysis (**Fig. 1c**)^26^. Proteins were quantified with the linear signal response expected from the dilution factors (**Fig. 1d**). Furthermore, quantitative reproducibility in replicates at the lowest level was still excellent (R = 0.96, **Fig. 1e**). Given that the protein amount of a single HeLa cell is as low as 150 pg^27^, and accounting for inevitable losses in sample preparation including protein digestion, we estimated that we would need to increase sensitivity by at least an order of magnitude to enable true single cell proteomics.

**Figure 1.**
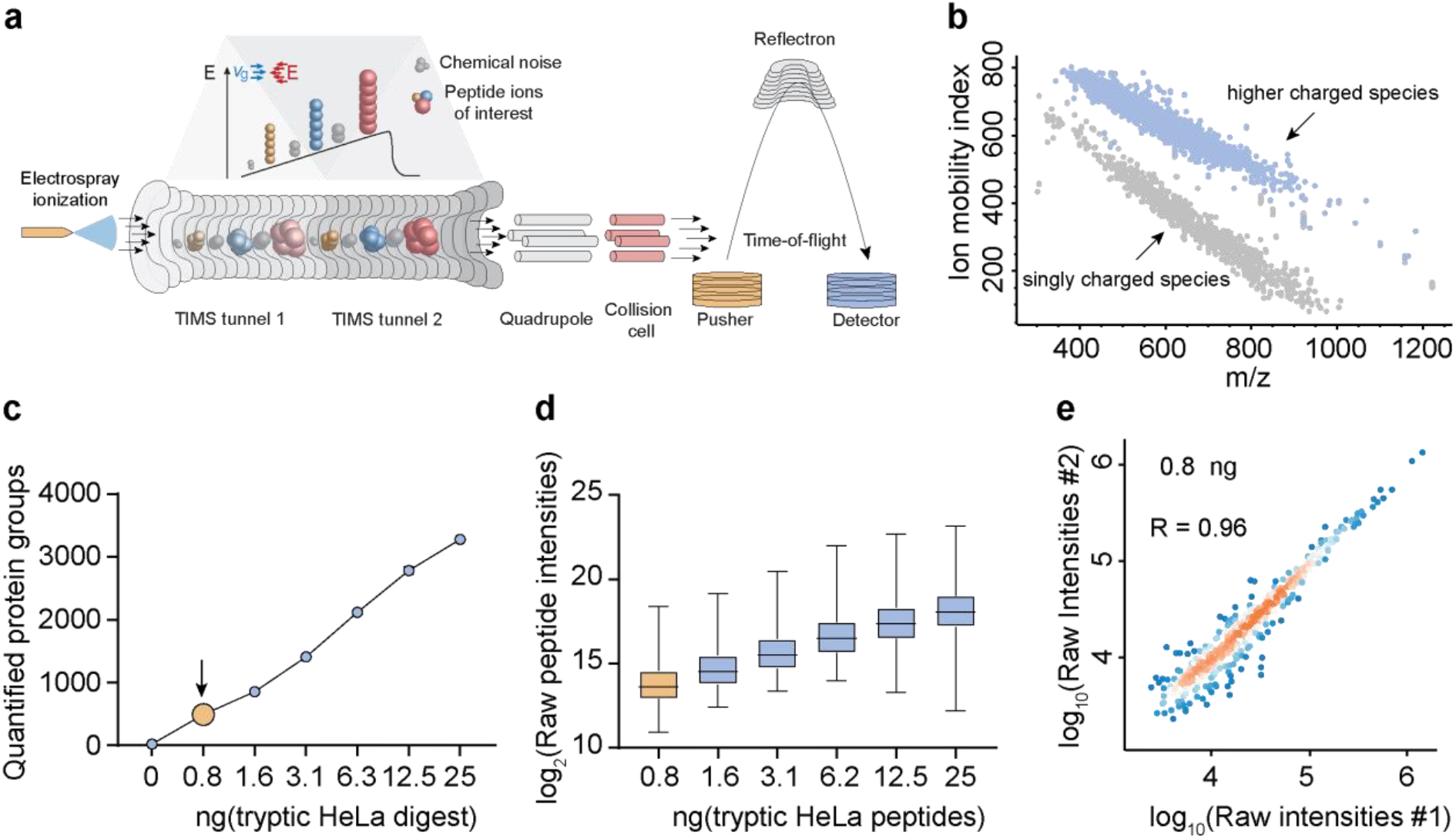
: TIMS enables virtually noise-free spectra and ultra-high sensitivity proteomics. **a, b**, TIMS-qTOF principle separating singly charged background peaks from multiply charged peptide precursor ions, making precursor ions visible at extremely low signal levels (0.8 ng HeLa digest). **c**, Quantified proteins from a HeLa digest dilution series from 25 ng peptide material down to 0.8 ng (arrow), roughly corresponding to the protein amount contained in three HeLa cells. **d**, Linear quantitative response curve of the HeLa digest experiment in c. **e**, Quantitative reproducibility of two successive HeLa digest experiments at the lowest dilution (technical LC-MS/MS replicates).

### True single-cell proteome analysis

Three main factors govern MS sensitivity: ionization efficiency, transfer efficiency into the vacuum system and ion utilization by the instrument^28^. We first constructed an instrument with a brighter ion source, introduced different ion optic elements and optimized parameters such as detector voltage. Together, this led to a more than 4-fold higher ion current (**Fig. 2a**). Next, we FACS sorted zero, one and up to six single HeLa cells in quadruplicate into individual 384-wells, processed them separately and analyzed them on this modified mass spectrometer. This resulted on average in 843, 1,279 and 1,890 identified proteins for one, two and six cells, respectively. Note that this analysis benefited from transferring peptide identifications on the MS1 level, as expected from extremely low sample amounts (**Fig. 2b**). Quantification accuracy was high when comparing single cells, not much reduced from comparing six cells (**Fig. 2 c, d**). A rank order abundance plot revealed that the measured single-cell proteome preferentially mapped to the higher abundant part of the six-cell proteome, indicating that proteome coverage depended deterministically on overall LC-MS sensitivity (**Fig. 2e**). Inspecting shared peptides between the single-cell and six-cell experiment showed that clearly interpretable precursor isotope patterns were still present at high signal-to-noise levels even at single-cell level following the cell count intensity ratio trend (**Fig. 2f**).

**Figure 2.**
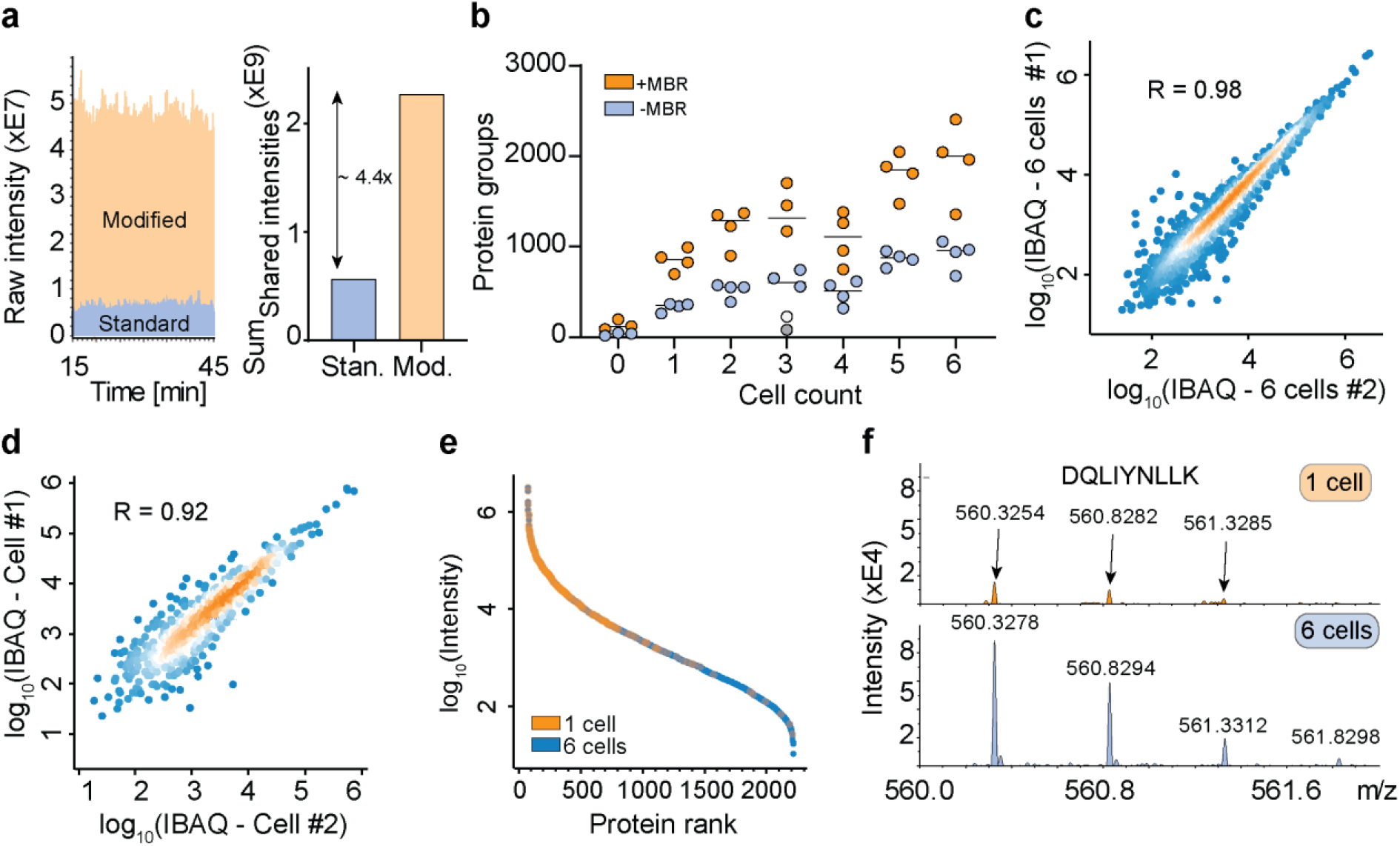
: A novel mass spectrometer allows the analysis of true single-cell proteomes. **a**, Raw signal increase from standard versus modified TIMS-qTOF instrument (left) and at the evidence level (quantified peptide features in MaxQuant) (right). **b**, Proteins quantified from one to six single HeLa cells, either with ‘matching between runs’ (MBR) in MaxQuant (orange) or without matching between runs (blue). The outlier in the three-cell measurement in grey (no MBR) or white (with MBR) is likely due to failure of FACS sorting as it identified a similar number of proteins as blank runs. **c**, Quantitative reproducibility in a rank order plot of a six-cell replicate experiment. **d**, Same as C for two independent single cells. **e**, Rank order of protein signals in the six-cell experiment (blue) with proteins quantified in a single cell colored in orange. **f**, Raw MS1-level spectrum of one precursor isotope pattern of the indicated sequence and shared between the single-cell (top) and six-cell experiments (bottom).

### More than 10-fold sensitivity increase

As electrospray (ES) is concentration dependent, sensitivity increases with decreasing flow-rate, however, very low flow systems are challenging to operate robustly and are consequently not widely available^28–30^. We recently described a chromatography system that decouples sample loading and gradient formation from the LC-MS run and operates at a standardized flow rate of 1 µL/min for high reproducibility^31^. This flow is fully controlled by a single pump instead of the binary gradients produced by other systems. We found that it worked robustly at flow rates down to 25 nL/min but standardized on 100 nL/min, which enabled stable operation for the entire project with the same column-emitter setup (**Extended data Fig. 1a, b**). ES sprayer diameter and gradient length were optimized for turnover, minimizing carry-over and stability.

MS-based T-SCP requires loss-less sample preparation by protein isolation and solubilization, followed by tryptic protein digestion and peptide purification ready for MS-analysis^20,21,32^. We found that small volumes of weak-organic solvents in conical 384-well plates provided a versatile and automatable environment for efficient cell lysis and protein digestion in minimal volumes (**Fig. 3a**). Briefly, single cells were sorted into wells containing 1 µL lysis buffer, followed by a heating step and further addition of buffer containing digestion enzymes to a total of 2 µL, all in an enclosed space. Peptides were concentrated in StageTip^33^ devices into 20 nL nanopackages, from which they were eluted in minimal volumes (**Fig. 3b**). To benchmark the effect of reduced flow rate and the concentrated peptide nanopackage elution, we directly compared signal traces of the normal 1 µL/min to the 100 nL/min set up. For 1 ng peptide material this resulted in a ten-fold increase in signal (**Fig. 3c**). To achieve high data completeness between hundreds of single-cell measurements, we next replaced ddaPASEF by diaPASEF, in which fragment level matching is further supported by ion mobility data^25^. We found that combining subsequent diaPASEF scan repetitions further improved protein identification numbers. Together, the very low flow chromatography and this diaPASEF acquisition mode resulted in the highly reproducible identification and quantification of more than 3,300 HeLa proteins from only 1 ng **(Fig. 3d)**, a drastic increase from the 550 identified in our initial set-up from a similar amount. Data completeness was at 94% and coefficient of variation (CV) less than 10 % for the selected scan repetition mode (**Extended data Fig. 1b**). This demonstrates that diaPASEF provides its advantages also at extremely low sample amounts, prompting us to adopt this acquisition mode for the single-cell workflow in the remainder of this work.

**Figure 3.**
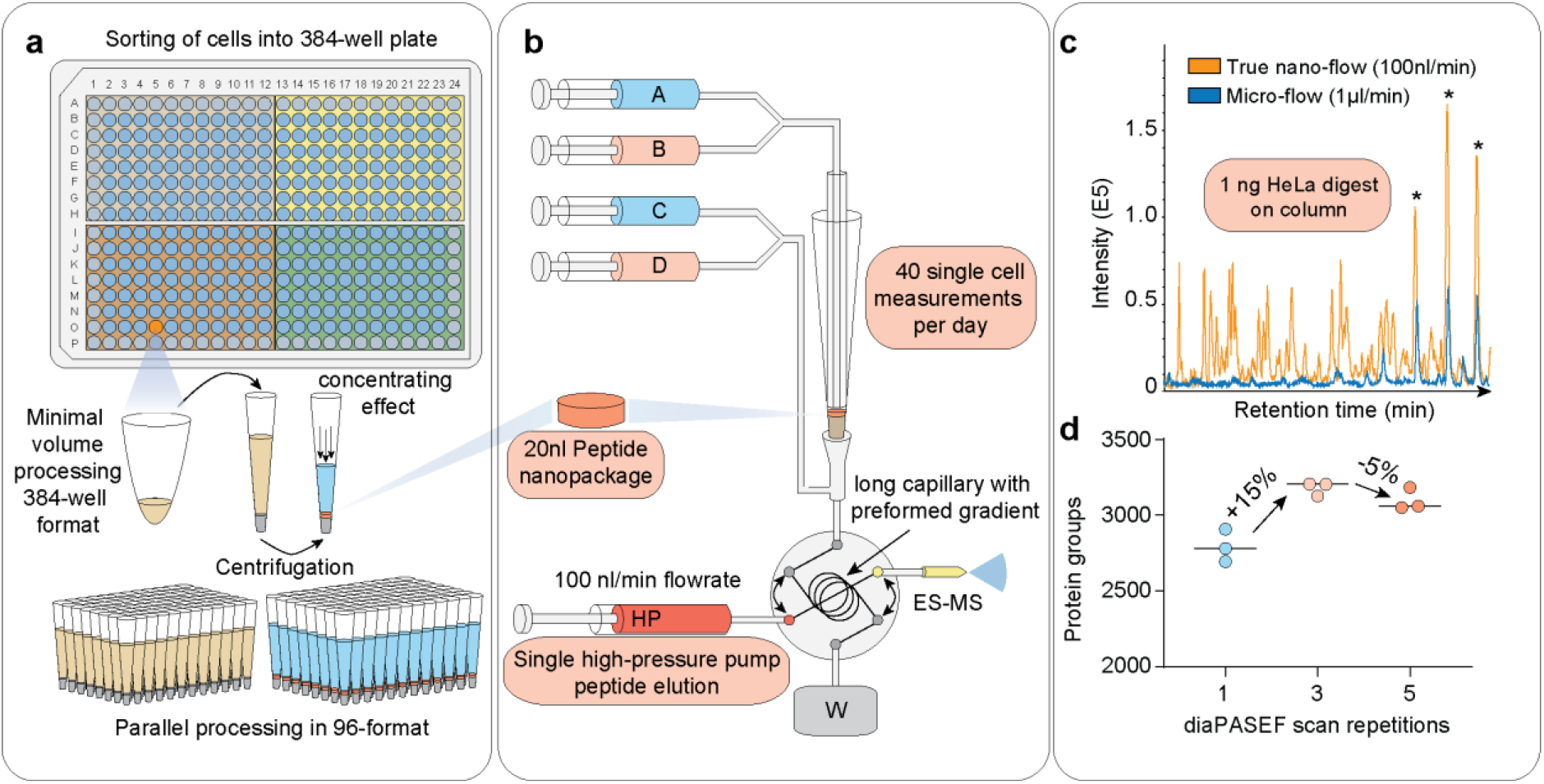
: Miniaturized sample preparation coupled to very low flow chromatography and diaPASEF. **a**, Single cells are sorted in a 384-well format into 1 µL lysis buffer by FACS with outer wells serving as qualitative and quantitaive controls. Single cells are lysed and proteins are solubilized at 72 °C in 20 % acetonitile, and digested at 37 °C. Peptides are concentrated into 20 nL nanopackages in StageTips in a 96-well format. **b**, These tips are automatically picked and peptide nanopackages are eluted in a sub-100 nL volume. After valve switching, the peptide nanopackage is pushed on the analytical column and separated fully controlled by the single high-pressure pump at 100 nL/min. **c**, Basepeak chromatogram of the standardized nano-flow (100 nL/min, orange) and micro-flow (1 µL/min, blue) gradients with 1 ng of HeLa digest on the StageTip. Asterices indicate polyethylene glycole contaminants in both runs. **d**, Nano-flow (100 nL/min) and short gradient diaPASEF method combined. Summation of 1 to 5 diaPASEF scan repetitions was used to find the optimum for high-sensitivity measurements at 1 ng of HeLa digest.

### T-SCP dissects arrested cell-cycle states

The cell cycle is an important and well-studied biological process that has frequently been used as a test case in single-cell studies^34,35^. To investigate if our proteomics workflow could detect biological responses to drug perturbation at the single-cell level, we treated HeLa cells with thymidine and nocodazole to produce four cell populations enriched in specific cell cycle stages (231 cells; **Fig. 4a**). We quantified up to 1,441 proteins per single-cell and 1,596 overall using a HeLa dia spectral library with about 34,000 precursors. This number ranged from a median of 611 in G1 to 1,263 in G1/S, 962 in G2, and 1,106 in G2/M (**Fig. 4b**). To estimate the total protein amount per cell, we summed all protein signals based on their identifying peptides. Judged by protein amount, G2 cells were approximately 1.5-fold larger than G1 cells; thus T-SCP correctly reflected the proliferation state, while highlighting a substantial heterogeneity within each that would have been hidden in bulk sample analysis (**Fig. 4c**). To be able to directly compare single-cell proteomes and cancel out protein abundance differences attributed to varying total protein amounts and identifications of each cell, we normalized our data set by local regression for all proteins with at least 15 % completeness across cells^36^. Furthermore, we stringently filtered our data set for at least 500 protein identifications per cell and more than 20% observations for each protein across remaining single cells (**Extended data Fig. 2a**). The proteomes of the different cell cycle states grouped together in a Principal Component Analysis (PCA) plot (**Fig. 4d**). In addition to these drug-perturbed cells, we measured more than 200 untreated ones from two independent cell culture batches to increase the overall numbers and representative proteome variability of single-cells measured. The T-SCP data set covered proteins assigned to many cellular compartments, membranes and biological processes involved in biological regulation, metabolism, transport, and signal transduction at high quantitative accuracy despite severe systematic perturbation introducing stark biological variation and proteome remodeling (**Extended data Fig. 2b, c; Extended data Table 1**). Next, we asked whether single-cell proteome measurements can be used to assign cellular states, similar to how single-cell RNA-sequencing (scRNA-seq) measurements have frequently been applied to cell type and state discovery, highlighted by cellular atlas projects^9^. In previous proteomics studies, cell populations had been enriched for cell cycle states and sets of regulated proteins had been extracted^34,35^. We here selected cell cycle stage marker proteins as the top 60 most differentially expressed in either the G2/M-, G1- and S-phase protein set from Geiger *et al*.^35^, as it used similar drug treatment on bulk populations and investigated how likely cells from different cell cycle stages could be distinguished (**Extended data Table 2**). We used these marker proteins to set up cell cycle stage specific scores indicating the likelihood to belong to the respective phase previously used for scRNA-seq cell cycle stage predictions. This model clearly distinguished cells from G2/M and G1/S and also other comparisons (**Fig. 4e, Extended data Fig. 2d**)^37^.

**Fig. 4:**
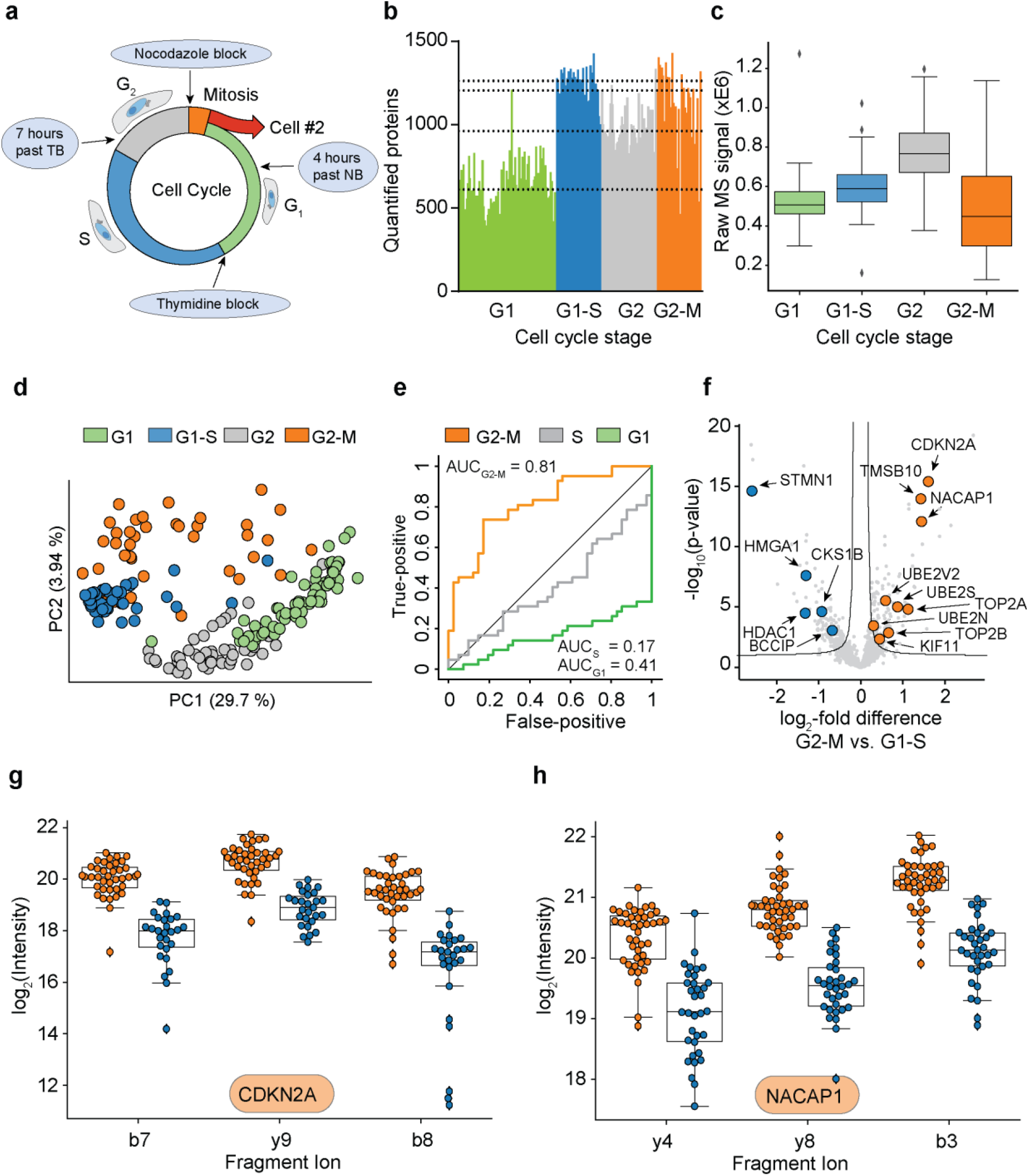
T-SCP correctly quantifies cell cycle states. **a**, Arresting single cells by drug perturbation. **b**, Numbers of protein identifications across 231 cells in the indicated cell cycle stages as enriched by the drug treatments in A. **c**, Boxplot of total protein signals of the single cells in B after filtering for at least 500 protein identifications per cell and 20% data completeness per protein across cells (G1: n = 69; G1-S: n = 41; G2: n = 51; G2-M: n = 42). **d**, PCA of single-cell proteomes of B. **e**, Receiver Operator Curve (ROC) for the prediction of G2/M cells against G1/S based on G2/M marker proteins with the indicated area under the curve (AUC) score. The other two curves, based on S and G1 marker proteins, respectively, indicate the inverse predictive power of these scores. **f**, Volcano plot of quantitative protein differences in the two drug arrested states. Arrows point towards colored significantly regulated key proteins of interest. **g**, Quantitative fragment ion level data for CDKN2A and its peptide ALLEAGALPNAPNSYGR (FDR < 10^−14^). **h**, Quantitative fragment ion level data for the pseudogene NACAP1 and its peptide IEDLSQEAQLAAAEK (FDR < 10^−14^).

Next, we investigated the differentially expressed proteins between the drug arrested cell cycle stage transition G2/M and G1/S. Among the significantly regulated proteins was a large number of known cell cycle regulators, some of which are highlighted (**Figure 4f; Extended data Table 3)**. Quantitative MS data at the fragment ion level was highly significant for these as illustrated by the cell cycle regulator CDKN2A and further examples (FDR < 10^−14^, **Fig. 4g**; **Extended data Fig. 3**). We also exemplary inspected precursor isotope patterns of CDKN2A on the raw MS1 and fragment ion series on the raw MS2 spectrum level highlighting the quality of ion signal and noise distribution by TIMS (**Extended data Fig. 4**). Our single-cell data set also highlighted proteins not previously associated with the cell cycle and the G2/M transition. For instance, the putative pseudogene NACAP1 was clearly identified and regulated (FDR < 10^−14^, **Fig. 4h, Extended data Fig. 5**). It might have escaped previous detection because of its small size (213 amino acids) as we have noticed previously^38^.

### SC proteomes compared to transcriptomes

Given our set of more than 430 single-cell proteomes, we compared the T-SCP measurements after filtering with similar single-cell RNA sequencing data (scRNA-seq)^39,40^. To achieve technology-independent insights, we selected assays from two widespread scRNA-seq technologies, Drop-seq^41^ and the lower throughput SMART-sSq2^42^, on the same cellular system. The Drop-seq assay is based on unique molecular identifiers (UMIs) to control for amplification biases in library preparation, whereas the SMART-Seq2 assay is not UMI-controlled. Note that MS-based proteomics inherently does not involve any amplification and is not subject to associated artifacts.

Despite subtle differences, HeLa cell culture should reflect a characteristic global distribution of gene and protein expression states^43^. This assumption would allow us to assess self-consistency of the measurement technologies. First, we computed the distribution over all pairwise correlation coefficients of cells within a technology^44^. We found that in the proteome measurement, cells have higher correlation on average than in the droplet-based method and similar correlation to the SMART-Seq2 method (**Extended data Fig. 6a**). On average in 54 % of the 1,596 proteins observed by MS-based proteomics, cells had non-zero expression values and the protein expression completeness per cell followed a normal distribution (**Fig. 5a**). For SMART-Seq2 this dropped to 28 % and to 7 % in the droplet-based protocol. Both single-cell RNA-sequencing technology data sets followed a bimodal gene completeness frequency distribution, while single-cell proteomes do not. (**Extended data Fig. 6b**). Next, we investigated whether there were fundamental limitations of the detection in the protein measurements. Such effects are discussed for scRNA-seq measurements as “drop-out events” or “zero-inflation”, although they are now much reduced in UMI-based protocols^44^. We identified signs of such detection limits as bimodality in the lower abundance range of the protein measurements (**Extended data Fig. 6c, d**), suggesting that our single-cell protein analysis could benefit from imputation or tailored likelihood-based parameter estimation methods^45^.

**Fig. 5:**
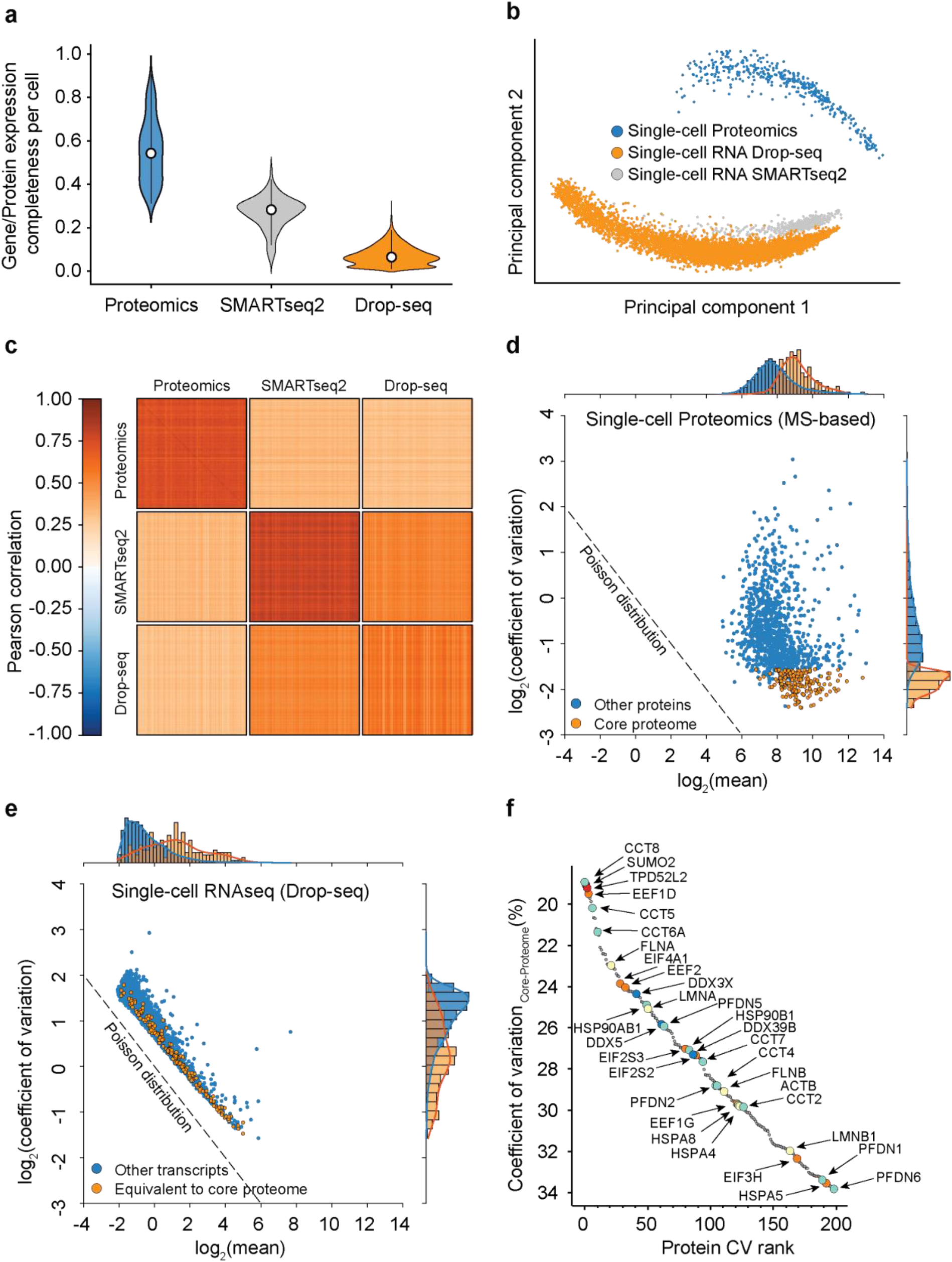
Single cells have a stable core proteome but not transcriptome. **a**, Gene or protein expression completeness per cell for either T-SCP (Cells x Proteins: 398 x 1,596), SMARTseq2 (Cells x Genes: 720 x 24,990), and Drop-seq (Cells x Genes: 5,383 x 41,161). **b**, Principal component analysis of single-cell gene and protein expression measurements. **c**, Heat map of cell-cell correlations across individual cells measured by proteomics and by both transcriptome technologies. **d**, Coefficient of variation of protein levels as a function of mean expression levels with the ‘core proteome’ colored in orange. **e**, Coefficient of variation of transcript expression values as a function of mean expression levels with transcripts corresponding to the core proteome in orange. **f**, Rank order abundance plot for the core proteome with color coded protein classes (Red: SUMO2 and TDP52L2 proteins; Turquoise: Chaperonin and folding machinery associated proteins. Orange: Translation initiation and elongation; Yellow: Structural proteins; Blue: DEAD box helicase family members).

For bulk measurements, transcript levels generally correlate moderately with the corresponding protein levels, however, this correlation strongly depends on the biological situation^46^. At the single-cell level this effect is further convoluted by technical differences in the measurement technologies. We asked to what degree scRNA-seq measurements could be used as a proxy for protein measurements in our data and found that protein measurements separate strongly from RNA in a principal component analysis (**Fig. 5b)**. Furthermore, single-cell transcript expression levels correlate well across scRNA-seq technologies, but not with single-cell protein measurements (**Fig. 5c, Extended data Fig. 6e**). This suggests that single-cell protein and RNA levels are very different, re-emphasizing that protein measurements yield complementary information to RNA measurements and do not simply re-iterate similar gene expression states. This implies distinct RNA and protein abundance regulation mechanisms on both modalities, dissection of which would not be possible with RNA measurements alone.

### T-SCP reveals a stable core proteome

Prompted by the divergent correlation values between the proteome and transcript levels, we next investigated the variability of gene expression as a function of abundance. For protein expression measurements, coefficients of variation were very small across covered abundances, independent of the data completeness (**Extended data Fig. 7a**). This is consistent with a model in which the covered proteome is stable and probed deterministically across its full dynamic range. In contrast, the same analysis for UMI-controlled and not UMI-controlled scRNAseq data revealed a much higher overall transcriptome variability, as measured by the coefficient of variation of single-cell RNA-seq compared to protein measurements (**Extended data Fig. 7a, c**). Remarkably, this difference is already very apparent with the current sensitivity of MS-based proteomics, which will surely increase in the future. Comparing single-cell proteome measurements with six-cell proteomes (**Fig. 2c**) suggests that a moderate increase in MS sensitivity would reveal a large part of the proteome to be quantitatively stably expressed. Based on these observations, we defined a ‘core-proteome’ subset in the MS-based proteomics data by selecting the top 200 proteins with the lowest CVs of the proteins shared between at least 70% of the more than 430 single-cells, including the drug perturbations (**Extended data Table 4**). Interestingly, these proteins where distributed well across the covered dynamic range of the proteome (**Fig. 6d**). Strikingly, we found the corresponding transcripts of the core proteome to be distributed across the full range of CVs in single-cell transcriptome data (**Fig. 6e, Extended data Fig. 7b, c**). The core proteome highlighted proteins frequently used for normalization such as HSP90 and ACTB, providing a positive control (**Fig. 5f**). The CV rank plot of the core proteome also reveals a diverse set of proteins, including representatives of translation initiation and elongation, folding machineries, nucleic acid helicases, as well as cellular structure determining proteins. Interestingly, we also identify TPD52L2 as one of the most stable proteins, which in turn is described as one of the most abundant proteins in HeLa cells^47^ and SUMO2, which is known for its involvement in a plethora of essential regulatory cellular processes, suggesting a stable cellular SUMO2-pool even during stark proteome remodelling^48^.

### Outlook

The T-SCP pipeline combines miniaturized sample preparation coupled to very low flow liquid chromatography and a novel mass spectrometer resulting in at least one order of magnitude sensitivity gain at highest robustness for the analysis of single cells. We quantify cellular heterogeneity following targeted perturbation, which enables the direct analyses of drug-responses in single-cell hierarchies on the proteome level. Furthermore, the comparison of single-cell RNA- and proteome level revealed that the proteome is stable while the transcriptome is more stochastic, highlighting substantial regulation of translation and setting the stage for its elucidation at the single cell level.

Although mainly demonstrated here for single-cell total proteome measurements, the sensitivity gain achieved in our workflow will be advantageous in any situation that is sample limited. This includes investigation of other compound classes such as metabolites or drugs, post-translational modifications from small numbers of cells or from *in vivo* material, measurements directly from paraffin embedded formalin fixed (FFPE) pathology specimens, which we are already pursuing^8^.

Our workflow is also compatible with chemical multiplexing with the advantage that the booster channel causing reporter ion distortions could be omitted or reduced. Furthermore, there are many opportunities for increasing overall sensitivity, including even brighter ion sources, improved chromatography and better data analysis and modeling tools, similar to the rapid recent advances in the scRNAseq field.

## Acknowledgements

A.-D.B. acknowledges support from the International Max Planck Research School for Life Sciences – IMPRS-LS. S.R. is supported by the Helmholtz Association under the joint research school “Munich School for Data Science” -MUDS. D.S.F. acknowledges support from a German Research Foundation (DFG) fellowship through the Graduate School of Quantitative Biosciences Munich (QBM). F.C. acknowledges the European Union’s Horizon 2020 research and innovation program (Marie Skłodowska-Curie individual fellowship under grant agreement 846795). We thank our colleagues in the Department of Proteomics and Signal Transduction, Max Planck Institute of Biochemistry, at the Center for Protein Research at Copenhagen University, and at Bruker Daltonik for discussions and help. In particular, we thank I. Paron, A. Piras and C. Deiml for technical support and J.B. Müller for column production. We also thank Mark Ridgeway, Craig Whitehouse, Andreas Brekenfeld, Niels Goedecke and Christoph Gebhardt of Bruker for their contribution to the development, installation and trouble-shooting of the modified mass spectrometer. We are grateful for the FACS-sorting support by the Imaging Core Facility at the Max Planck Institute of Biochemistry, in particular Martin Spitaler and Markus Oster.

## Author contributions

A.-D. B. and M.M. conceptualized and designed the study. A.-D.B., M.C.T., F.C., and A.M. performed experiments. M.A.P. and O.R. designed the new mass spectrometer. A.-D.B., O.B.H. and N.B. conceived the new EvoSep gradient. O.B.H., N.B., A.-D.B. and M.C.T. designed the new EvoSep gradient and optimized it for proteomics performance. D.F. and F.J.T. conceptualized the single cell modeling. A.-D.B., S. R., D. F., F.J.T., C. A., M.C.T., O.B.H., N.B., C.V., F.M. and M.M. analyzed the data; A.-D.B. and M.M. wrote the manuscript.

## Methods

### Sample preparation for bulk dilution experiments

For all benchmark experiments purified peptides from bulk HeLa cells were used. HeLa was cultured in Dulbecco’s modified Eagle’s medium at 10 % fetal bovine serum, 20 mM glutamine and 1% penicillin–streptomycin. Cells were collected by centrifugation, washed with phosphate-buffered saline (PBS), flash-frozen in liquid nitrogen and stored at −80 °C. Cells were resuspended in PreOmics lysis buffer (PreOmics GmbH) and boiled for 20 min at 95 °C, 1500 rpm to denature, reduce and alkylate cysteins, followed by sonication in a Branson, cooled down to room temperature and diluted 1:1 with 100 mM TrisHCl pH 8.5. Protein concentration was estimated by Nanodrop measurement and 500 µg were further processed for overnight digestion by adding LysC and trypsin in a 1:50 ratio (µg of enzyme to µg of protein) at 37 °C and 1500 rpm. Peptides were acidified by adding 1 % trifluoroacetic acid (TFA) 99 % isopropanol (IprOH)in a 1:1 ratio, vortexed, and subjected to StageTip^1^ clean-up via styrenedivinylbenzene reverse phase sulfonate (SDB-RPS). 20 µg of peptides were loaded on two 14-gauge StageTip plugs. Peptides were washed two times with 200 µL 1 % TFA 99 % IprOH followed by 200 µL 1 % TFA 99 % IprOH in an in-house-made StageTip centrifuge at 2000 xg and elution with 100 µL of 1 % Ammonia, 80 % acetonitrile (ACN), 19% ddH2O into PCR tubes and finally dried at 60 °C in a SpeedVac centrifuge (Eppendorf, Concentrator plus). Peptides were resuspended in 0.1 % TFA, 2 % ACN, 97.9 % ddH_2_O.

### Sample preparation for single-cell experiments

HeLa cells were cultured as described above. Supernatant was removed, cells were detached with trypsin-treatment, followed by strong pipetting for cell aggregate dissociation. Cells were washed three times with ice-cold Phosphate-buffered Saline (PBS), pelleted by centrifugation, and the supernatant was removed. For fluorescent-activated cell-sorting (FACS), DAPI was added and sorting performed on the DAPI-negative live cell population. Single cells were sorted into 384-well plates containing 1µl of 20 % acetonitrile (ACN), 100 mM TrisHCl pH 8.5, centrifuged briefly, sealed with aluminum foil and frozen at −80 °C until further use. When needed, single-cell containing 384-well plates were incubated for 30 min at 72 °C in a PCR cycler, followed by 5 min sonication (Elmasonic P) at 37 kHz and room temperature. Protein digestion was performed overnight at 37 °C in a PCR cycler after adding 1 µL of 20 % ACN, 100 mM TrisHCl pH 8.5, 1 ng trypsin/lysC mix. For the peptide bulk and cell count dilution experiments, peptides were resuspended in 4 µL of 2 % ACN, 0.1 % TFA, 97.9 % ddH_2_O and injected directly via NanoLC. For all other single cell experiments, samples were dried in a SpeedVac, resuspended in 20 µL 0.1 % formic acid (FA), 99.9 % ddH2O (Buffer A) before transfer into activated EvoTips. These were activated following the standard EvoSep protocol^2^. Then, 50 µL buffer A was added to each EvoTip followed by centrifugation at 200 xg for 1 min. The sample was transferred into the EvoTip, followed by centrifugation at 600 xg for 1min, and two centrifugation steps after adding 50 µL buffer A. Last, 150 µL buffer A was added to each EvoTip and spun for 30 sec at 300 xg.

### Cell cycle experiments

The drug-perturbed cell cycle arrest experiment was designed to enrich cells in four cell-cycle stages – G1, the G1/S-transition, G2 and the G2/M-transition. HeLa cells were grown to approximately 30 % confluence as described above, washed and treated for 24 h with 5 mM thymidine, released for 4.5 h and treated again with 5 mM thymidine, or 0.1 µg/mL nocodazole for 13 h. Cells of the G1/S phase (thymidine block) or G2/M phase (nocodazole block) were washed in PBS, trypsinated, subjected to strong pipetting to dissociate cell aggregates and ice-cold PBS washes before DAPI-negative single live cell FACS sorting. A second set of G1/S phase and G2/M phase blocked cells was washed and cultured for 7 h or 2.5 h to enrich early G2 and G1-phase HeLa cells. These were washed with PBS, trypsinated and subjected to DAPI-negative single live cell FACS sorting into 384-well plates pre-loaded with 1 µL 20 % acetonitrile, 100 mM TrisHCl pH 8.5 lysis buffer. Furthermore, we prepared presumable unsynchronized cells sets from two independent cell cultures and subjected them to sample preparation as described below.

### High-pH reversed-phase fractionation

To generate a deep library of HeLa precursors for all data-dependent benchmark experiments, peptides were fractionated at pH 10 with the spider-fractionator^3^. 50 μg of purified peptides were separated on a 30 cm C_18_ column in 96 min and concatenated into 24 fractions with 2 min exit valve switches. Peptide fractions were dried in a SpeedVac and reconstituted in 2 % ACN, 0.1 % TFA, 97.9 % ddH_2_O for LC-MS analysis.

### Liquid-chromatography

For the initial benchmark experiments with HeLa bulk dilution and the cell count dilution, liquid chromatography analysis was performed with an EASY nanoLC 1200 (Thermo Fisher Scientific). Peptides were loaded on a 45 cm in-house packed HPLC-column (75 µm inner diameter packed with 1.9 µm ReproSil-Pur C18-AQ silica beads, Dr. Maisch GmbH, Germany). Sample analytes were separated using a linear 60 min gradient from 5-30 % B in 47.5 min followed by an increase to 60 % for 2.5 min, by a 5 min wash at 95 % buffer B at 300 nL/min and re-equilibration for 5 min at 5 % buffer B (Buffer A: 0.1 % Formic Acid (FA), 99.9 % ddH2O; Buffer B: 0.1 % FA, 80 % ACN, 19.9 % ddH2O). The column temperature was kept at 60 °C by an in-house manufactured oven.

For all other proteome analyses, we used an EvoSep One liquid chromatography system^4^ and analyzed the single-cell proteomes with a novel 35 min stepped pre-formed gradient eluting the peptides at 100 nL/min flow-rate. We used a 15 cm × 75 μm ID column with 1.9 μm C18 beads (EvoSep) and a 10 µm ID electrospray emitter (Bruker Daltonik). Mobile phases A and B were 0.1 % FA in water and 0.1 % FA in ACN, respectively.

Both LC systems were coupled online to a modified trapped ion mobility spectrometry quadrupole time-of-flight mass spectrometer (timsTOF Pro, Bruker Daltonik GmbH, Germany) via a nano-electrospray ion source (Captive spray, Bruker Daltonik GmbH).

### Construction of a novel mass spectrometer with higher sensitivity

We modified our ion source to draw more ions into the vacuum system of the instrument by modifying the glass capillary that conducts gas and ions between the ionization region at atmospheric pressure and the first pumping region of the mass spectrometer. Additional gas is eliminated via an extra pumping stage. Novel prototype ion optics, a high-pressure ion funnel and a radio frequency (RF) multipole confine the ions and transport them to the next vacuum region where the analysis by trapped ion mobility mass spectrometry (TIMS) occurs. The glass capillary is oriented orthogonal to the high-pressure funnel so that neutral contaminants and droplets are first directed away from the funnel by the gas flow. Furthermore, the high-pressure funnel and RF multipole are oriented orthogonal to the TIMS, maintaining the gas dynamics of our original design. Remaining neutral contaminants are guided away from the TIMS entrance. To accommodate the increased ion current, the TIMS analyzer was updated to a new stacked ring (SRIG) design. We use a higher order RF field in the ion accumulation region to create a larger effective ion storage volume than the low order fields of previous designs. A low order quadrupolar field is maintained in the analyzer region to compress the ions towards the analyzer axis during elution to maintain high mobility resolution. The transition between the high order and low order parts of the device was optimized compared to prior designs to further improve peak shape and ion mobility resolution. This results in about a factor of three gain in ion capacity and therefore about a factor of three in the instrument’s dynamic range.

### Mass spectrometry

Mass spectrometric analysis was performed either in a data-dependent (dda) or data-independent (dia) PASEF mode. For ddaPASEF, 1 MS1 survey TIMS-MS and 10 PASEF MS/MS scans were acquired per acquisition cycle. Ion accumulation and ramp time in the dual TIMS analyzer was set to either 50/100/200 ms each and we analyzed the ion mobility range from 1/K_0_ = 1.6 Vs cm^-2^ to 0.6 Vs cm^-2^. Precursor ions for MS/MS analysis were isolated with a 2 Th window for m/z < 700 and 3 Th for m/z >700 in a total m/z range of 100-1,700 by synchronizing quadrupole switching events with the precursor elution profile from the TIMS device. The collision energy was lowered linearly as a function of increasing mobility starting from 59 eV at 1/K_0_ = 1.6 VS cm^-2^ to 20 eV at 1/K_0_ = 0.6 Vs cm^-2^. Singly charged precursor ions were excluded with a polygon filter (otof control, Bruker Daltonik GmbH). Precursors for MS/MS were picked at an intensity threshold of 1.500 arbitrary units (a.u.) and re-sequenced until reaching a ‘target value’ of 20.000 a.u. considering a dynamic exclusion of 40 s elution. For DIA analysis, we made use of the correlation of Ion Mobility (IM) with m/z and synchronized the elution of precursors from each IM scan with the quadrupole isolation window. We used the described 100ms ddaPASEF method for the acquisition of a HeLa bulk single-shot library for the single-cell experiments and the short gradient diaPASEF method as described in Meier *et al* ^5^, but performed up to 5 consecutive diaPASEF cycles before the next MS1-scan (see main text). The collision energy was ramped linearly as a function of the IM from 59 eV at 1/K0 = 1.6 Vs cm^−2^ to 20 eV at 1/K0 = 0.6 Vs cm^−2^.

### Raw data analysis

ddaPASEF data for tryptic HeLa digest dilution series and the cell count experiment were analyzed in the MaxQuant environment (version 1.6.7) and searched against the human Uniprot databases (UP000005640_9606.fa, UP000005640_9606_additional.fa), which extracts features from four-dimensional isotope patterns and associated MS/MS spectra^6,7^. False-discovery rates were controlled at 1% both on peptide spectral match (PSM) and protein levels. Peptides with a minimum length of seven amino acids were considered for the search including N-terminal acetylation and methionine oxidation as variable modifications and cysteine carbamidomethylation as fixed modification, while limiting the maximum peptide mass to 4,600 Da. Enzyme specificity was set to trypsin cleaving c-terminal to arginine and lysine. A maximum of two missed cleavages were allowed. Maximum precursor and fragment ion mass tolerance were searched as default for TIMS-DDA data. Peptide identifications by MS/MS were transferred by matching four-dimensional isotope patterns between the runs (MBR) with a 0.7-min retention-time match window and a 0.05 1/K_0_ ion mobility window in case of the single cell-count dilution experiment into a deep ddaPASEF library consisting of 24 fractionations of tryptic HeLa digest. These data were also searched without matching between runs to access the MBR-mediated identification increase. Either intensity-based absolute quantification (IBAQ) or label-free quantification was performed with the MaxLFQ algorithm and a minimum ratio count of one^8^.

For all other single-cell experiments, we used a small precursor library consisting of 34,195 precursors mapped to 28,235 peptides and 4,536 protein groups, which was acquired with the 100ms ddaPASEF method described above and generated with the Spectronaut software (version 14.10.201222.47784; Biognosys AG, Schlieren, Switzerland)^9^. A minimum of three fragments per peptide, and a maximum of six fragments were included. All single-cell measurements were searched against the human UniProt reference proteome (UP000005640_9606.fa, UP000005640_9606_additional.fa) of canonical and isoform sequences. Searches used protein N-terminal acetylation and methionine oxidation as variable modifications. We generated one decoy precursor per precursor in the spectral library and used a conservative normal distribution estimator approach for p-value estimation. Protein intensities were normalized using the “Local Normalization” (Q-value = 0.15) algorithm based on a local regression model^10^. A protein and precursor FDR of 1% was used. Default settings were used for other parameters.

### Visualization and FDR estimates of fragment ion intensities

Quantitative fragment ion profiles were generated from the Spectronaut output table via the “F.PeakArea” column. Only fragment ions used for quantification in Spectronaut were included (EG.UsedForProteinGroupQuantity = True, EG.UsedForPeptideQuantity = True, F.ExcludedFromQuantification = False). To cancel out cell-size dependent abundance changes, one normalisation factor was estimated per cell, using fold-change based normalization of the whole dataset, as described in the MS-EmpiRe method, which we also used for FDR control^11^. The intensities were log2 transformed and subsequently visualized.

### Proteomics downstream data analysis

Proteomics data analysis was performed in the Perseus environment (version 1.6.7, 1.5.5)^12^, GraphpadPrism (version 8.2.1) and Python (version 3.8.2). MaxQuant output tables were filtered for ‘Reverse’, ‘Only identified by site modification’, and ‘Potential contaminants’ before further processing. Ontologies for the biological process and cellular compartment assignment for proteins was performed with the mainAnnot.homo_sapiens.txt.gz followed by categorical counting across all proteins for each of the ontologies and counts were exemplary visualized as frequency plot. For single-cell analysis, if not otherwise specified, the Spectronaut data output was filtered first for at least 500 protein observations per cell and at least 20% quantification events across rows and log(x+1)-transformed. The single-cell run with the analysis number 306 was removed since it was identified as full quantitative outlier via PCA analysis. For correlation analysis of two protein expression vectors, transformed gene or protein quantification events of two cells were plotted against each other replacing missing values by zeros. For principal component analysis (PCA), missing values were imputed from a normal distribution with a width of 0.3 standard deviations that was downshifted by 1.8 standard deviations. Differential expression analysis by two-sided unpaired t-test was performed on two groups filtered for at least 50% row-wise quantification events within one group. False-discovery rate control due to multiple hypothesis testing was performed by a permutation-based model and SAM-statistic with an S_0_-parameter of 0.3. For cell-size estimation based on raw MS-signal, intensity outputs within cell cycle resolved single-cell proteomics results were summed up and visualized as boxplots. The core proteome was calculated by filtering the whole single-cell proteomics data set for at least 70% quantification events for each protein followed by selection of the top 200 proteins with the smallest coefficient of variation across the dataset.

### Single-cell protein and RNA comparison and dropout statistics

The SMART-Seq2^13^ data set measured 720 HeLa cells in 3 different batches with a total of 24,990 expressed genes. The Drop-seq^14^ data set contained 3 batches with a total of 5,665 cells and 41,161 expressed genes. We performed the single cell analysis with scanpy v1.6.0^15^. If not stated otherwise, we used standardized filtering across all datasets, removed cells with less than 500 genes expressed and removed genes detected in less than 20% of the remaining cells, resulting in 9,900 transcripts in 720 cells in the SMART-Seq2 dataset and 5,000 transcripts and 5,383 cells measured with Drop-seq technology. Ratios of non-zero entries in the scRNAseq datasets and the number of identified proteins in our data are summarized as violin plots. To investigate data completeness across covered dynamic range, we computed the data completeness as a function of the mean log(x+1)-transformed protein abundance of all non-zero/-NaN entries. We included the expected data completeness based on the assumption that missing values are purely due to shot-(Poisson)-noise as 1-exp(-x). For correlation analysis, the RNA abundance entries were linearly scaled to sum to the mean cell size of the respective dataset per cell (230,881.33 for SMART-Seq2 and 6,948.1 for Drop-Seq) followed by log(x+1)-transformation of all abundance entries. Correlation values between the expressions of two cells were computed as the Pearson correlation on the 1073 genes that were shared in all 3 datasets. Entries of missing protein abundance values were excluded from the specific computation. For the PCA plot of technological comparisons, the gene coverage intersection of all technologies (1,073) was isolated, NaNs were replaced by zeros, and expression values were linearly scaled to 1E6 followed by log(1+x) transformation. In coefficient of variation (CV) versus CV plots comparing different technologies as well as the mean versus CV analysis (including the core proteome analysis) and the CV distribution boxplots, RNA expression vectors were scaled to the mean cell size of that measurement technology and mean and CV values were computed per gene under the assumption that single-cell RNA-sequencing data are not zero-inflated^16^. CV (Proteomics) versus CV (RNA-seq) plots show the comparison of cv values of proteins/genes that were shared between all datasets.

### Cell Cycle State Prediction

Cell cycle predictions were performed using the scanpy method score_genes^15^ based on three sets of proteins that are specifically expressed in the G1-(MARCKS, LMO7, KRT1, GDA, KRT2, HIST1H1E, KRT18, HNRNPA1, DBNL, OGT, CHCHD3, CD44, DBN1, NASP, TARDBP, SH3BGRL3, PODXL, SUMO2, ZYX, STMN1, BAG3, TRIM28, PGRMC1, COASY, EFHD2, SPTAN1), S-(NOLC1, ATP2A2, CANX, CPT1A, TMX1, CKB, SLC25A3, ATP1B3, SLC16A1, MT-CO2, EPHX1, SRPRB, CYB5R3, TECR, LETM1, ANP32B, NUP205), or G2/M-phase (TOP2A, TOP2B, HMGB1, EIF5B, TMSB10, NCAPD2, EIF3D, ANP32A, SELENBP1, BAZ1B, RCC2, S100A4, FASN, HINT1, DKC1, LUC7L2, AARS, KPNA2, CKAP5), respectively. The cell phase specific protein sets were selected based on the z-scored fold-change ratios provided in Geiger *et al*.^17^. The top60 highest differentially expressed genes were selected, but only the aforementioned ones were also identified in our single-cell proteomics data. This scoring method yields the average expression on the provided set of genes minus the average expression on a reference set of genes for each cell. The reference set is chosen to mirror the average expression of the target gene set. For this analysis, cells and genes were filtered, log(x+1)-transformed and missing values replaced by zeros. Plotted are the ROC curves for the three scores corresponding to the three sets of characteristic proteins (G1, S and G2M) used individually to discriminate between the cells of two cell cycle stages.

### Data availability

All mass spectrometry raw data, libraries and outputs from each particular search engine analyzed in this study have been deposited to the ProteomeXchange Consortium via the PRIDEpartner repository and are made available to the reviewers.

## Extended data Figures

**Extended data Figure 1:**
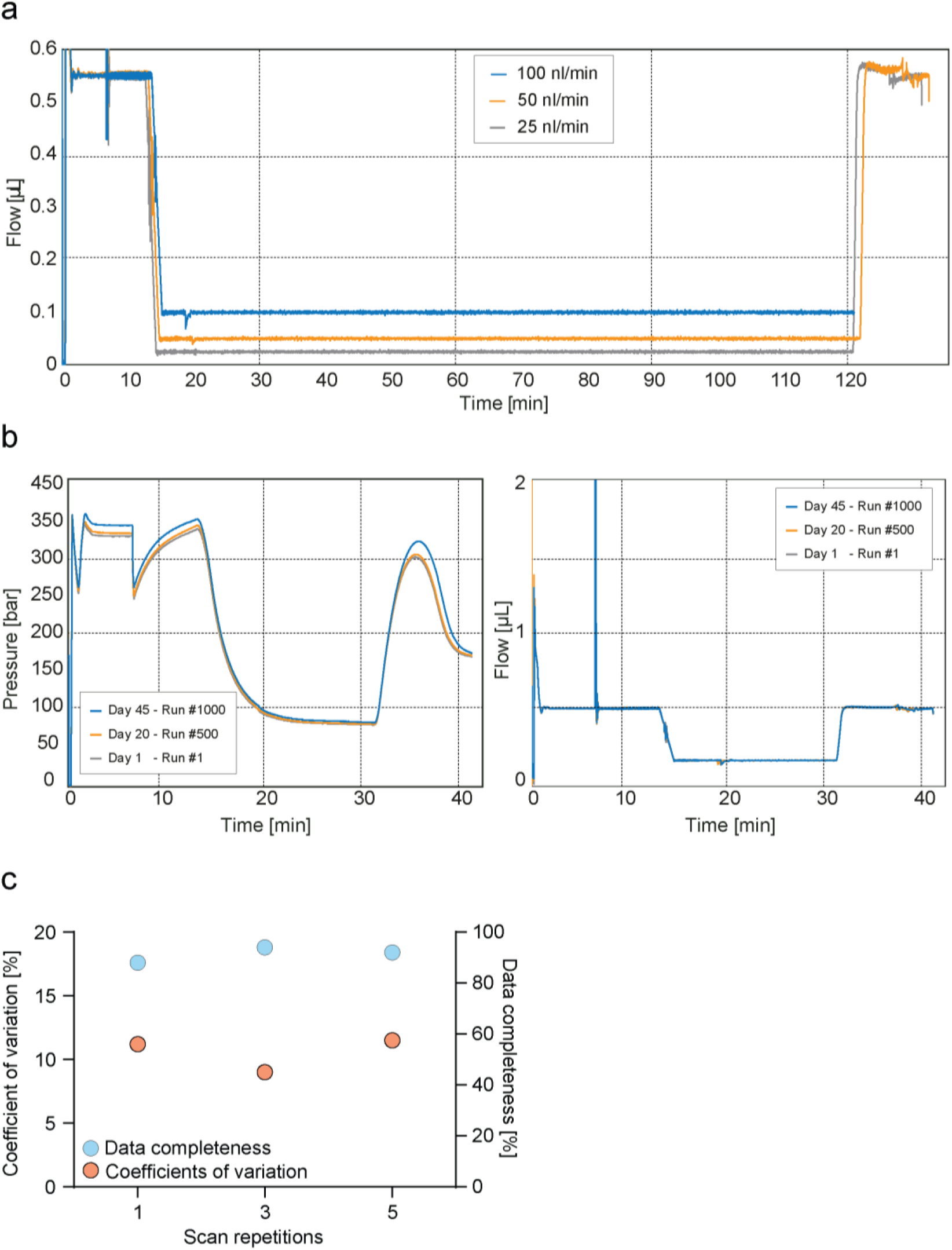
Robust very low flow rate chromatography and performance evaluation of diaPASEF for ultra-high sensitivity proteomics. **a**, True nanoflow at 25, 50, and 100 nl/min flow rate on the EvoSep One liquid chromatography system. **b**, Standardized 100 nl/min true nano flow gradient on the EvoSep One liquid chromatography system. Pressure (Left) and flow profile (right) of the gradient of more than 1000 consecutive runs (Day 1 – Run #1 = gray; Day 20 – Run #500 = orange; Day 45 – Run #1,000 = blue). **c**, Data completeness (Blue) and coefficient of variation (Orange) evaluation of different diaPASEF consecutive scan repetitions merged for the analysis of 1 ng tryptic HeLa digest. Scans were varied from one, three, and five repetitions.

**Extended data Figure 2:**
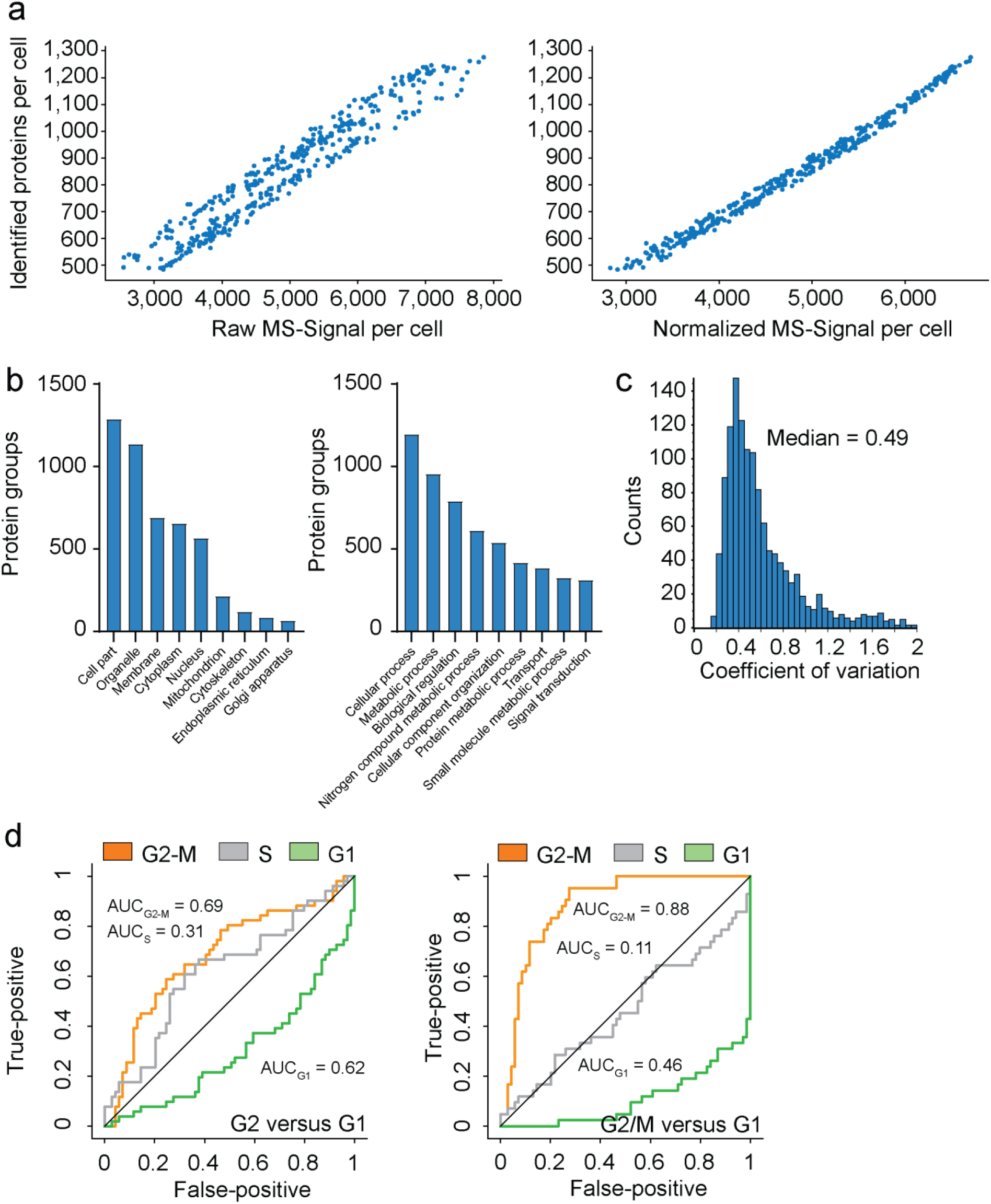
Qualitative and quantitative investigation of T-SCP data. **a**, Raw log(x+1)-transformed intensity values of proteins per cell plotted against the number of identified proteins per cell (Left) and after normalization by local regression to cancel out those differences to enable downstream analysis (Right). **b**, Category count of gene ontology annotations for cellular compartment and biological process terms. Exemplary, category count terms are shown for the cellular compartment (left) and biological process (right) for the more than 430 single-cell proteomics data set. **c**, Frequency plot for coefficient of variation occurrence within the 420 single cell proteomics data set. **d**, Cell cycle stage prediction for G2 versus G1 phase cells (left) and G2/M versus G1 phase cells (right) using the top 60 most differentially expressed proteins reported by Geiger *et al*. as input.

**Extended data Figure 3:**
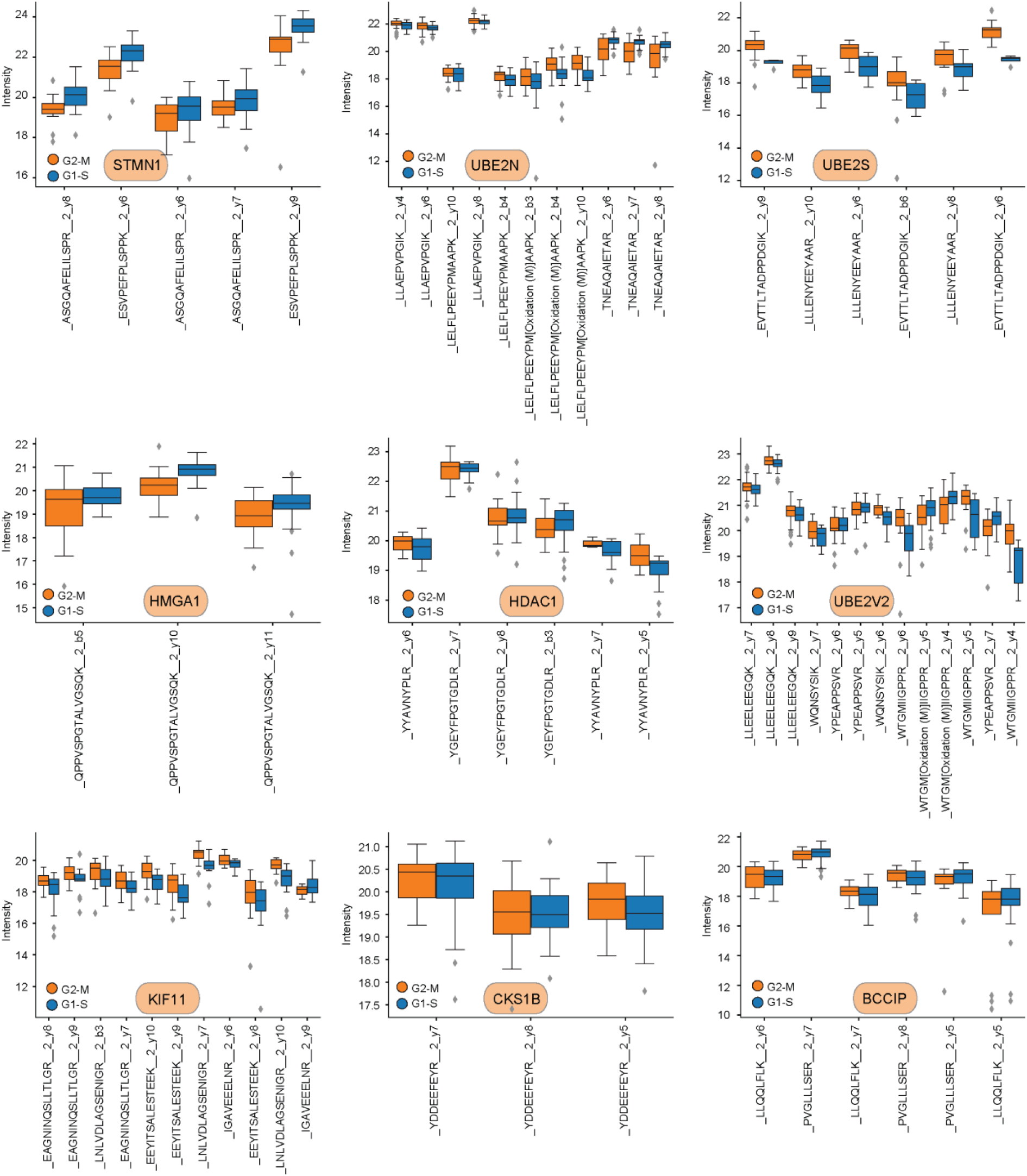
Fragment ion intensities of peptides for several differentially expressed proteins (UBE2S, FDR = 2.96E-10; STMN1, FDR = 0.00159; HMGA1, FDR = 0.00664; BCCIP, FDR = 0.92523; HDAC1, FDR = 0.83649; UBE2V2, FDR = 0.5152; KIF11, FDR = 2.38E-6; CKS1B, FDR = 0.60734; UBE2N, FDR = 0.7521) in the comparison of nocodazole (G2-M transition) and thymidine (G1-S transition) treated cells. Boxplots represent the intensity distribution of indicated peptide fragment ion intensities (G2-M: red; G1-S: blue).

**Extended data Figure 4:**
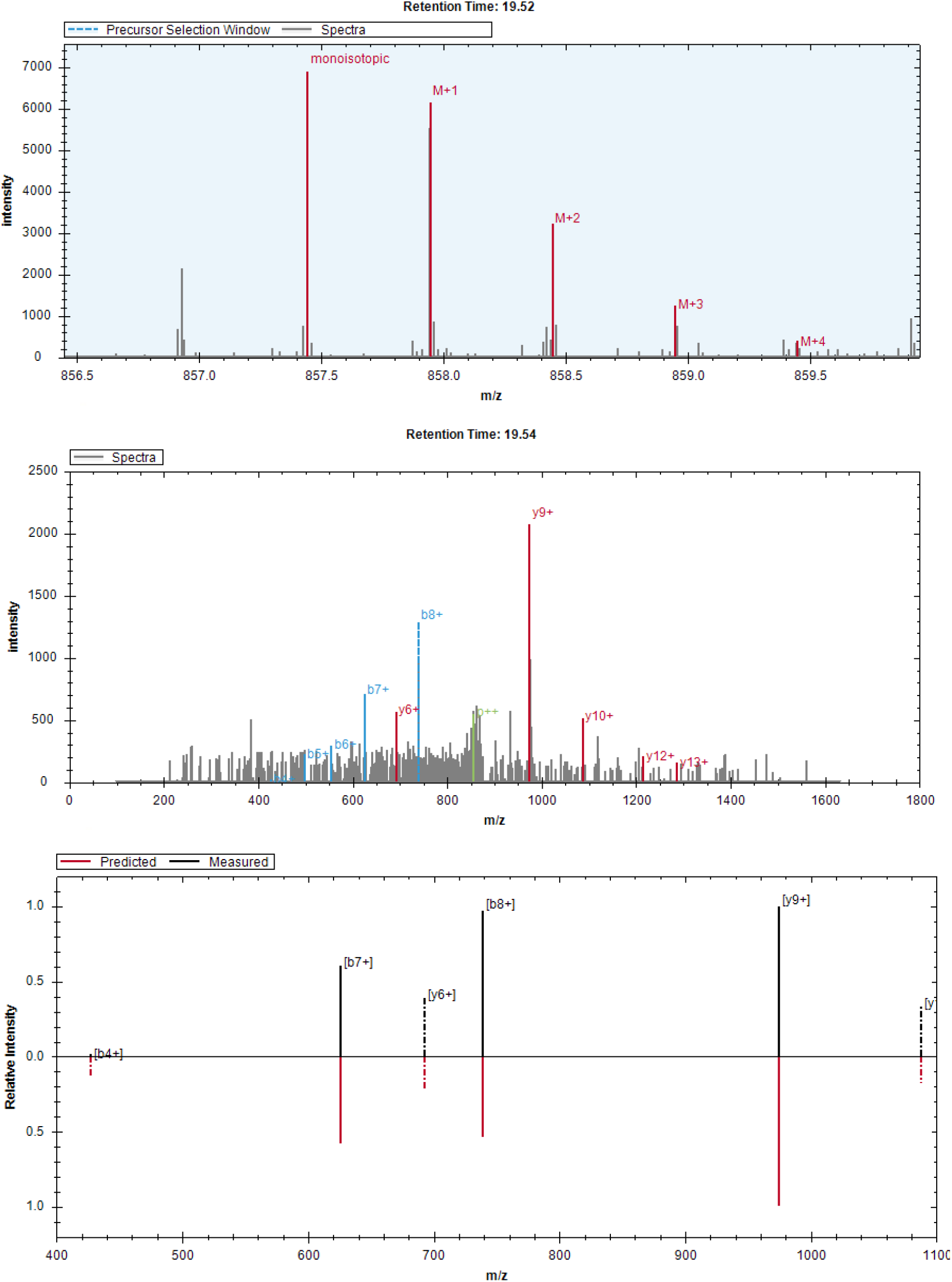
Exemplary raw MS1 isotope pattern level for the CDKN2A peptide ALLEAGALPNAPNSYGR at MS1 spectrum elution apex (Retention time: 19.52 min; top) and raw MS2 b-(blue) and y-(red) fragment ion series at elution apex (p++ = Unfragmented precursor-ion signal; Retention time: 19.54 min; middle). MS2 intensity correlation of predicted and experimental fragment ion series and intensity values of the same peptide (bottom).

**Extended data Figure 5:**
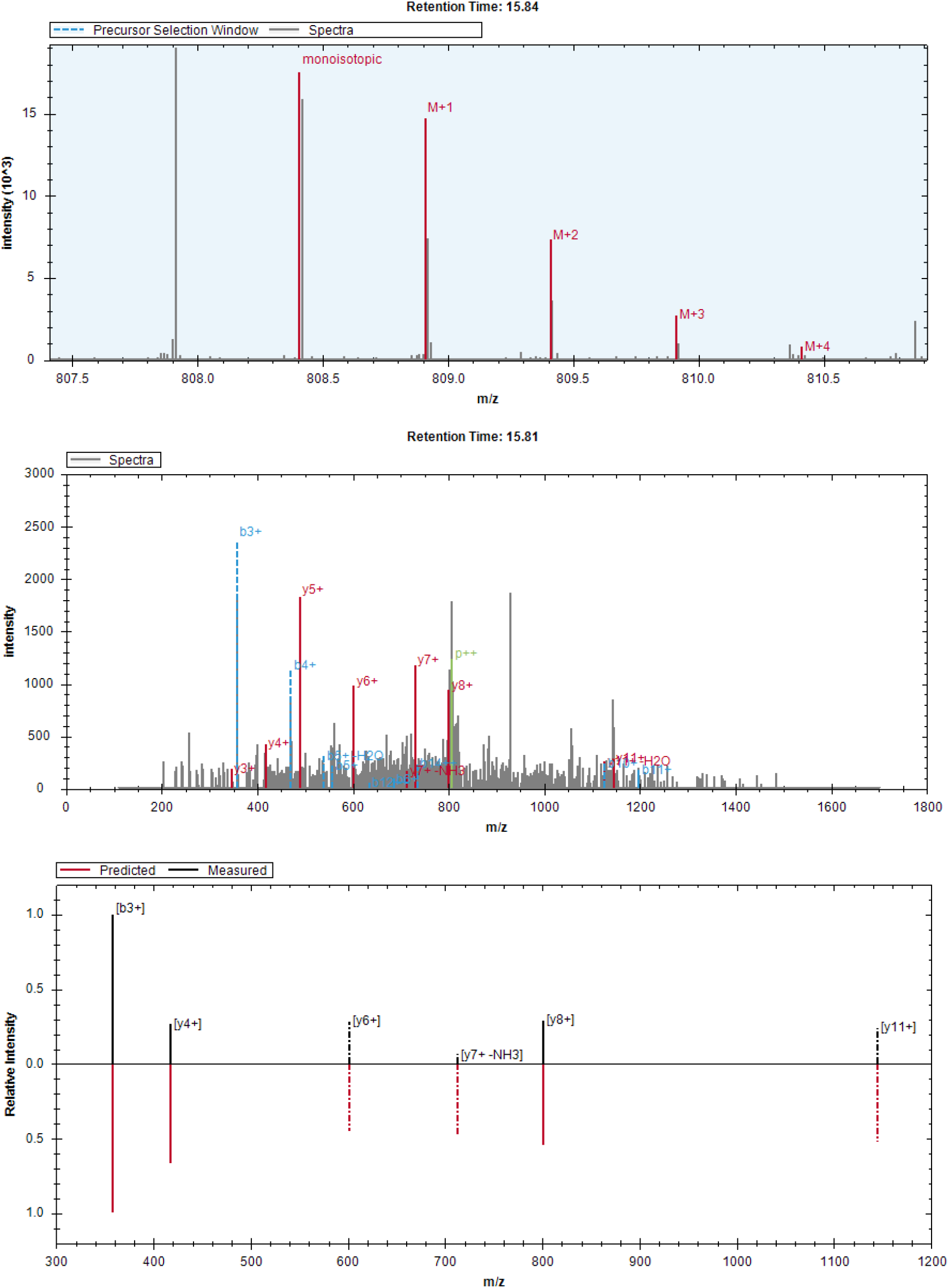
Exemplary raw MS1 isotope pattern level for the NACAP1 peptide IEDLSQEAQLAAAEK at MS1 spectrum elution apex (Retention time: 15.84 min; top) and raw MS2 b-(blue) and y-(red) fragment ion series at elution apex (p++ = Unfragmented precursor-ion signal; Retention time: 15.81 min; middle). MS2 intensity correlation of predicted and experimental fragment ion series and intensity values of the same peptide (bottom).

**Extended data Figure 6:**
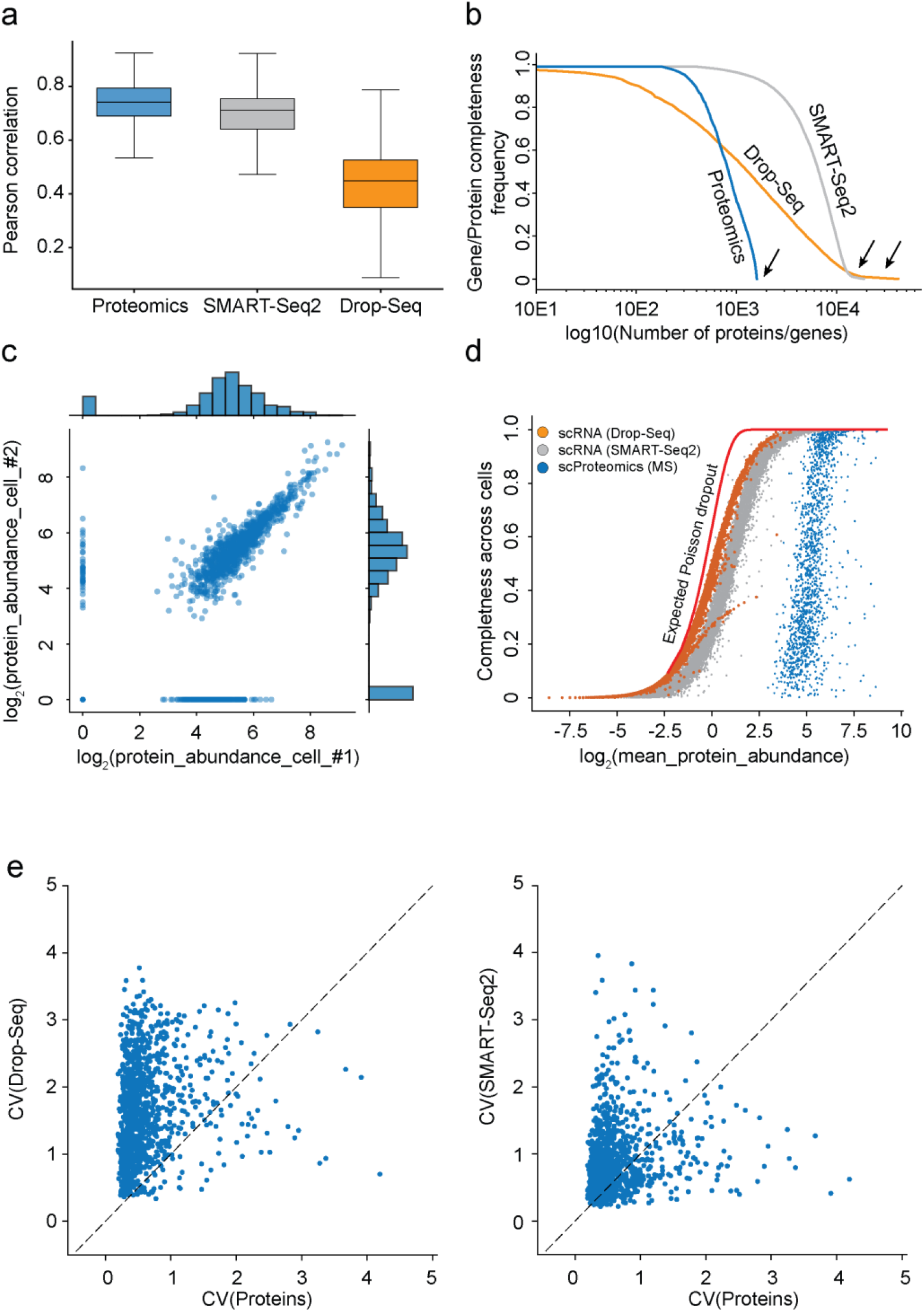
Comparison of single-cell RNA-sequencing and proteomics data. **a**, Pearson correlation of observations for each cell within each of the technologies (MS-based proteomics, SMART-Seq2 RNA-sequencing, droplet-based RNA-sequencing). **b**, Gene or protein expression frequency occurrence as a function of completeness higher than X for all three technologies (Proteomics, dropSEQ, SMART-Seq2). Arrows indicate a bimodal distribution for single-cell RNAseq data in both technologies, while absent in Proteomics. **c**, Scatter plot of two independently measured single-cell proteome expression values. **d**, Data completeness across single cells as a function of mean protein abundance for MS-based single-cell proteomics and both single-cell RNA-sequencing (Drop-Seq, SMART-Seq2). Expected poison dropout distribution shown in red. **e**, The coefficient of variation of a gene measured by either Drop-Seq technology (left) or SMART-Seq2 (right) compared to the coefficient of variation of the corresponding protein measured by MS-based single-cell proteomics.

**Extended data Figure 7:**
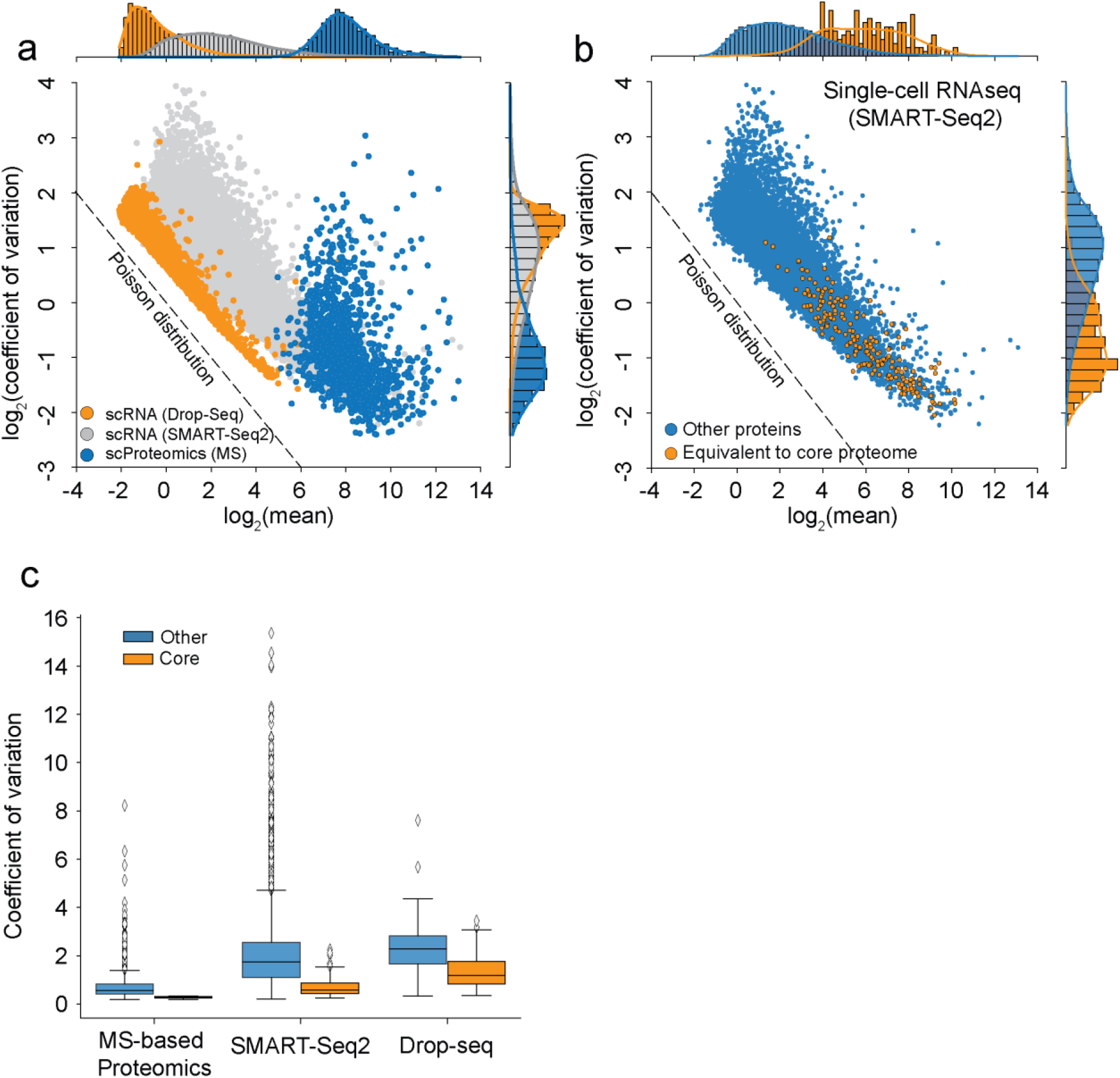
Single cells have a stable core proteome, but not transcriptome. **a**, Coefficient of variation distribution as a function of mean gene or protein intensities for either Drop-Seq (orange), SMART-Seq2 (gray), or MS-based single-cell proteomics (blue). Expected Poisson distribution shown as dashed line. **b**, Coefficient of variation of single-cell RNA-sequencing (SMART-Seq2) levels as a function of mean expression levels with the ‘core proteome’ colored in orange and non-’core proteome’ genes in blue. Expected Poisson distribution shown as dashed line. **c**, Coefficients of variation for each protein or gene expressed across the data set for either MS-based single-cell proteomics, SMART-Seq2 and Drop-Seq dataset. Proteins or genes of the ‘core-proteome’ are shown in orange, others in blue.

